# Cluster dispersal shapes microbial diversity during community assembly

**DOI:** 10.1101/2025.04.17.649399

**Authors:** Loïc Marrec, Sonja Lehtinen

**Affiliations:** Département de Biologie Computationnelle, Université de Lausanne, Lausanne, Switzerland; Swiss Institute of Bioinformatics, Lausanne, Switzerland

## Abstract

Identifying the drivers of diversity remains a central challenge in microbial ecology. In microbiota, within-community diversity is often linked to host health, which makes it all the more important to understand. Since many communities assemble de novo, microbial dispersal plays a critical role in shaping community structure. While theoretical models typically assume microbes disperse individually, this overlooks cases where microbes disperse in clusters, such as, for example, during host feeding. Here, we investigate how cluster dispersal impacts microbial community assembly, species richness, between-community dissimilarity, and species abundance. We developed a model in which microbes disperse from a pool into communities as clusters and then replicate locally. Using both analytical and numerical approaches, we show that cluster dispersal promotes community homogenization by increasing within-community richness and reducing dissimilarity across communities, even at low dispersal rates. Moreover, it modulates the influence of local selection on microbial community assembly and, consequently, on species abundance. Our results demonstrate that cluster dispersal has distinct effects from simply increasing the dispersal rate. We discuss how these predictions can inform the interpretation of experimental data and demonstrate their utility by reanalyzing a gut microbiota dataset from *Caenorhabditis elegans*. This analysis reveals new evidence for the role of cluster dispersal in microbial community assembly.

## Introduction

Dispersal is a key driver of microbial community assembly (Clobert et al., 2001). Many microbial communities form from scratch when new microenvironments emerge. For example, many organisms do not inherit their parents’ microbiota and are therefore born with germ-free microbiota, which become populated after birth [e.g., *Caenorhabditis elegans* (Zhang et al., 2017), *Drosophila melanogaster* (Blum et al., 2013)]. The early dispersing microbes are crucial, as they shape community composition, metabolic activity, and the development of complex ecosystems and biogeochemical processes (Hadland et al., 2024). Thus, community assembly is a potentially critical phase in the development of microbial diversity.

In the context of host-associated microbiota, within-community microbial diversity is often linked to host health. However, the nature of this relationship varies depending on the body site. For instance, a healthy vaginal microbiota is typically characterized by low species richness (Baud et al., 2023). Its composition becomes especially crucial during pregnancy, where it serves as a barrier against infections that could threaten both maternal and fetal health (Baud et al., 2023). Conversely, in the gut, reduced microbial diversity is frequently associated with a range of adverse health outcomes (Larsen and Claassen, 2018), including inflammatory, metabolic, and immune-related disorders. Interventions aimed at increasing gut microbiota diversity, such as the use of prebiotics, probiotics, or fecal microbiota transplantation, have shown therapeutic potential in restoring microbial diversity and improving host health (van Nood et al., 2013; Tillisch et al., 2013).

Microbial diversity encompasses more than just within-community variation. Microbial diversity also includes substantial differences between communities, often referred to as between-community dissimilarity (or *β*-diversity). Microbial communities can vary dramatically in composition and abundance across individuals, even among hosts of the same species (Spor et al., 2011; Lundberg et al., 2012; Smith et al., 2015). Notably, even genetically identical individuals, such as monozygotic twins, often harbor markedly different microbiomes, highlighting the role of non-genetic influences in shaping microbial ecosystems (Goodrich et al., 2014; Vilchez-Vargas et al., 2022).

Dispersal plays a key role in within- and between-community diversity. Limited dispersal increases between-community dissimilarity, while high dispersal rates homogenize community structures (Etienne and Olff, 2004). Vega and Gore (2017) experimentally confirmed this by assembling worm gut microbiotas with two species, demonstrating that richness increases with dispersal, whereas between-community dissimilarity decreases. They quantified this transition using the bimodality coefficient (Ellison, 1987), a summary statistic describing abundance fluctuation distributions (Grilli, 2020), which is the distribution of abundances across communities that are replicates of a same assembly processes.

Building on Vega and Gore (2017)’s work, Marrec and Bank (2024) developed a community assembly model to derive analytical predictions for the bimodality coefficient, refining assessments of dispersal’s role in richness and between-community diversity. They also introduced mean relative abundance, i.e., the extent to which the community is dominated by a single species, as an additional metric for comparing microbial traits across species.

Like many theoretical models (e.g., Zapién-Campos et al. (2020)), Marrec and Bank (2024) assumed that microbes disperse individually. However, in host-associated communities, microbes are often ingested in clusters. For example, humans regularly ingest multiple microbes through food and water. Thus, a single meal often introduces a mix of multiple microbes into the human gut. Obadia et al. (2017) experimentally measured microbial establishment probabilities in the fruit fly *D. melanogaster* gut using inoculum doses from 10^1^ to 10^8^ CFUs. Similarly, Jones et al. (2022) used a 5 × 10^6^ CFU inoculum. To date, it remains unclear how cluster dispersal impacts the assembly dynamics of microbial communities.

In this study, we investigate how cluster dispersal influences richness and between-community dissimilarity. We develop a model in which two microbial species disperse from a pool into local communities, in which they replicate. Using both analytical and numerical approaches, we demonstrate that cluster dispersal tends to homogenize microbial communities and influences mean relative abundance in a non-monotonic way when combined with within-community selection. We then apply our results to Vega and Gore (2017)’s experimental data, shed light on cluster dispersal, and even estimate cluster size. Finally, to assess the robustness of our predictions, we extend our model to multiple species and quantify *α*- and *β*-diversity. Overall, our work highlights the role of cluster dispersal in microbial community assembly.

## Model and methods

### Microbial community assembly model

We build a model to represent the assembly of microbial communities that are initially microbe-free. This model, shown in Figure 1A, includes a microbial pool consisting of two species, A and B, present in abundances *p*_A_ and 1 − *p*_A_, respectively. Microbial clusters of size *n* disperse from the pool into local communities at a rate *c*. The composition and abundance of these clusters are drawn from a binomial distribution ℬ (*n, p*_A_). Note that the local communities are assumed to be isogenic, meaning that selection is homogeneous across them. This assumption is relevant to certain experimental setups involving clonal worms or flies (Vega and Gore, 2017; Ortiz et al., 2021; Jones et al., 2022).

**Figure 1:**
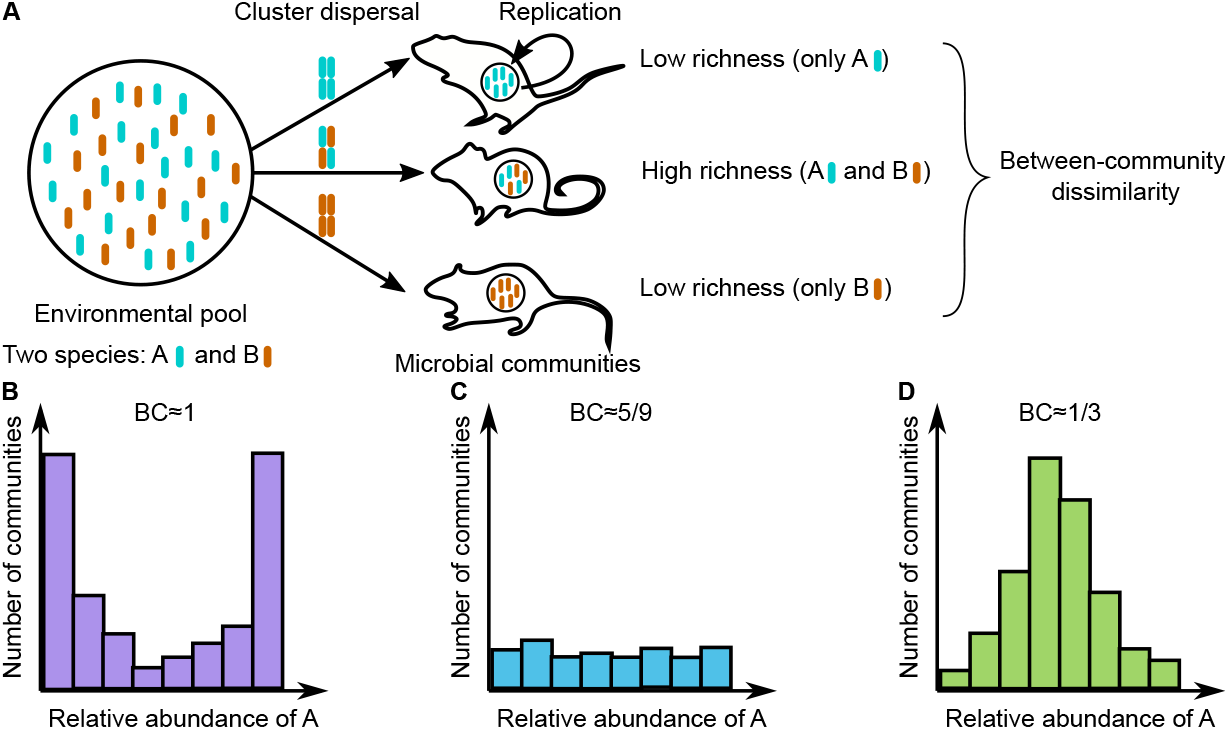
Sketch of the microbial community assembly model. As shown in Panel **A**, the environmental pool contains two species, A (blue) and B (orange). Species A is present in the pool in abundance *p*_A_, while species B is present in abundance 1 − *p*_A_. Microbial clusters of size *n* disperse from the pool into local microbial communities at rate *c* and replicate within these communities at rates *r*_A_ and *r*_B_. Cluster composition and abundance are drawn from a binomial distribution ℬ (*n, p*_A_). Once each community reaches its carrying capacity, we analyze the abundance fluctuation distribution, which quantifies the number of communities with a given composition and abundance (**B, C**, and **D**). We characterize this distribution using its bimodality coefficient (BC) and mean relative abundance.

Once introduced into a community, microbes replicate at rates that may depend on the species: *r*_A_ for species A and *r*_B_ for species B. To account for the limiting effects of carrying capacity *K* on community growth, the rates are multiplied by a saturation term, namely (1 − *N/K*), which is derived from logistic growth dynamics (Tsoularis and Wallace, 2002). Here, *N* = *N*_A_ + *N*_B_ represents the total community size. We consider that the assembly process is complete when the community size reaches the carrying capacity (*N* = *K*).

Once community assembly is complete, we quantify species richness and between-community dissimilarity (i.e., within- and between-community diversity, or *α*- and *β*-diversity, respectively) using the abundance fluctuation distribution (Grilli, 2020), which describes the number of communities that contain a given abundance of A microbes (Figures 1B, C, and D). To characterize this distribution, we use the bimodality coefficient (BC), a summary statistic of probability distributions that ranges from 0 to 1 (Ellison, 1987). Values of BC exceeding 5/9 indicate bimodality, suggesting a low-dispersal assembly regime dominated by cell replication (*c* ≪ *r*; Figure 1B). This regime typically leads to low richness but high between-community dissimilarity (Vega and Gore, 2017; Marrec and Bank, 2024). In contrast, BC values below 5/9 signify a unimodal distribution, characteristic of a high-dispersal regime where dispersal events dominate (*c* ≫ *r*; Figure 1D), resulting in high richness and low between-community dissimilarity (Vega and Gore, 2017; Marrec and Bank, 2024).

An additional descriptor of the abundance fluctuation distribution is its mean, which can be used to compare microbial traits across species and to detect processes such as selection (Marrec and Bank, 2024).

### Microbial community assembly simulation

We simulate the assembly of a microbial community using a Gillespie algorithm (Gillespie, 1976, 1977), which generates trajectories of stochastic dynamics based on known event rates. In our model, three types of events are considered: the replication of microbes from species A and species B

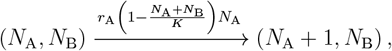

and

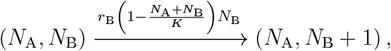

and the dispersal of clusters

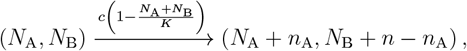

where *n*_A_ is drawn from a binomial distribution ℬ (*n, p*_A_). For each event, the propensity function is indicated above the arrow. The simulation steps are as follows:

1. Initialization: The microbial community starts from *N*_A_ = 0 and *N*_B_ = 0 microbe.
2. Event selection: The next event to occur is chosen randomly proportionally to its probability. For example, the replication of a microbe of species A is chosen with probability *r*_A_*N*_A_*/*(*r*_A_*N*_A_ + *r*_B_*N*_B_ + *c*).
3. Population size update: The population sizes *N*_A_ and *N*_B_ are updated according to the event selected in Step 3. For example, if the replication of a species A microbe is chosen, the population size of species A is updated by *N*_A_ ← *N*_A_ + 1.
4. We return to Step 2 until the community size is equal to the carrying capacity (*N*_A_ +*N*_B_ = *K*).

Each stochastic realization of the above algorithm describes the assembly of a single microbial community. Thus, collecting several stochastic realizations is equivalent to simulating the microbial assembly of several communities.

## Data availability

Simulations were performed with Matlab (version R2024b). All annotated code to reproduce the simulations and visualizations is available at https://github.com/LcMrc and will be deposited on Zenodo upon acceptance of the paper.

## Results

### Cluster dispersal homogenizes microbial communities

To examine the impact of cluster dispersal on microbial community assembly, we first consider the neutral case, where both species share the same replication rate (*r*_A_ = *r*_B_ = *r*). Specifically, we simulate community assembly across various dispersal rates while keeping the replication rate constant and quantify the bimodality coefficient BC of the resulting abundance fluctuation distributions (Figures 1B, C, and D).

We derive the expression for the bimodality coefficient under these two regimes (Supporting Information - Section 1)

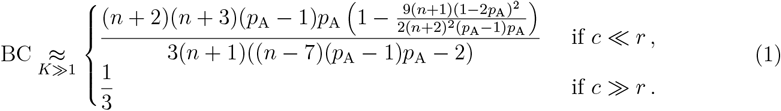

Assuming that both species are present in equal abundance in the microbial pool (*p*_A_ = 1*/*2) simplifies the previous equations to

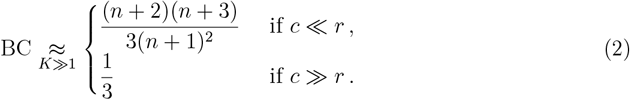

These equations show that, in the low-dispersal regime, the bimodality coefficient BC ranges from 1 when *n* = 1 to 1/3 when *n* ≫ 1, which is identical to the BC value observed in the high dispersal regime. Consequently, large clusters yield the same bimodality coefficient in both assembly regimes, making them indistinguishable through this metric.

Figure 2A shows the bimodality coefficient (BC) as a function of the dispersal rate. As expected, BC approaches 1/3 at high dispersal rates, indicating a unimodal abundance fluctuation distribution, characteristic of high richness and low between-community dissimilarity. This behavior is independent of cluster size, as dispersal in this regime homogenizes community composition to reflect that of the microbial pool.

**Figure 2:**
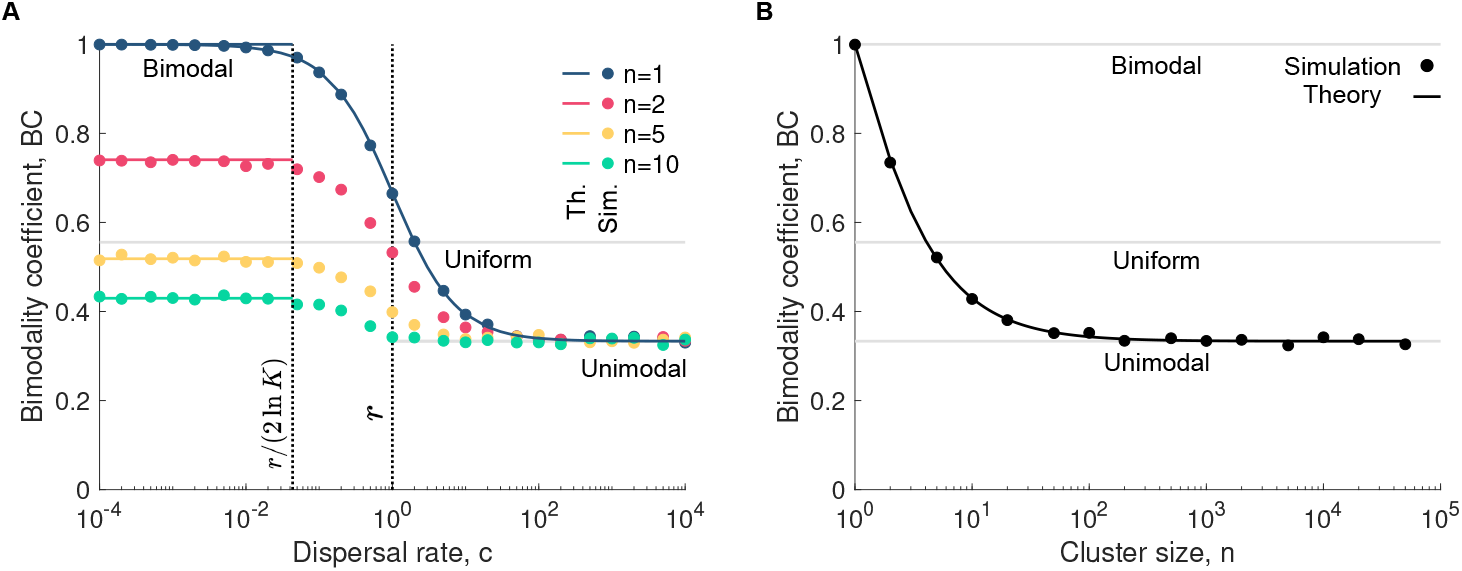
Cluster dispersal blurs the boundary between assembly regimes. Panel **A** shows the bimodality coefficient BC as a function of the dispersal rate *c* for various cluster sizes *n*, whereas panel **B** shows it as a function of the cluster size *n*. In both panels, the simulated data are averaged over 10^4^ microbial communities. The solid lines represent our analytical predictions (see Equation 2). Parameter values: replication rates *r*_A_ = *r*_B_ = 1, dispersal rate *c* = 10^−4^ (in **B**), relative abundance of A in the pool *p*_A_ = 1*/*2, carrying capacity *K* = 10^5^.

In contrast, in the low-dispersal regime, the bimodality coefficient depends strongly on cluster size, as shown in Figure 2B. When cluster size is small (e.g., 1), communities are typically populated by a single species, leading to a BC of 1, signifying low richness and high between-community dissimilarity. As cluster size increases, the probability of introducing multiple species in the initial dispersal event rises. This increases richness, decreases between-community dissimilarity and, thus, reduces the bimodality coefficient toward 1/3. For example, if *p*_A_ = 1*/*2, the probability that a cluster of size 2 contains only one species is 1/4, while for size 10, it drops to 1/1024. Therefore, larger clusters promote more diverse initial dispersal events, shifting community assembly dynamics toward those observed in the high-dispersal regime.

It is important to note that accounting for the dispersal of clusters is not equivalent to increasing the dispersal rate of individual microbes. Even when the bimodality coefficient is plotted as a function of the effective dispersal rate, that is, the number of individual microbes dispersing per unit time rather than the number of clusters, its maximum value still decreases with increasing cluster size (Supporting Information - Figure S2). This is because larger clusters are more likely to introduce multiple species simultaneously.

Equation 1 and Figure 2B together provide a useful framework for determining whether a given cluster size homogenizes microbial communities and supports distinguishable assembly regimes.

### Cluster dispersal modulates the impact of local selection on microbial community assembly

A second metric for characterizing microbial community assembly is the mean relative abundance. The mean relative abundance quantifies the extent to which a community is dominated by a single species. Thus, this metric can be used to determine, for example, if local selection is present, i.e., whether two species have different replication rates (Marrec and Bank, 2024). Whereas in the previous section we focused on the neutral case (*r*_A_ = *r*_B_), we now assume that the two species have different replication rates (*r*_A_ ≠ *r*_B_), resulting in a nonzero selection coefficient *s* = *r*_A_ − *r*_B_, and investigate the impact of within-community selection on the mean relative abundance. Note that incorporating within-community selection has little impact on the bimodality coefficient, which remains similar to that obtained in the neutral case (Supporting Information - Figure S4).

Analytically, we build on Houchmandzadeh (2018)’s work and derive an equation giving the mean absolute abundance of A once the community assembly is complete, denoted by ⟨*N*_A_⟩. Note that here the mean relative abundance is simply given by the mean absolute abundance of A divided by the carrying capacity once the community assembly is complete. In the low-dispersal regime, we obtain the following expression (Supporting Information - Section 1)

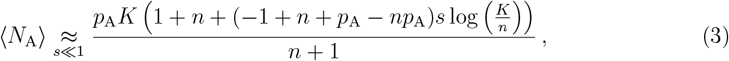

which, assuming that both species are present in equal abundance in the pool (*p*_A_ = 1*/*2), reduces to

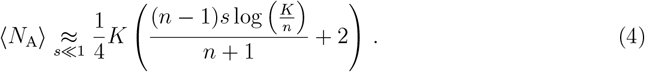

If the selection coefficient is zero, the mean absolute abundance is equal to *p*_A_*K* for any cluster size. This value is also obtained for a cluster size of 1 or *K* for any selection coefficient. If the selection coefficient is nonzero, the mean absolute abundance varies non-monotonically with increasing cluster size, reaching a maximum that exceeds 50% when *s >* 0 and a minimum below 50% when *s <* 0. In the high-dispersal regime, the community structure is expected to reflect that of the microbial pool such that the mean absolute abundance satisfies ⟨*N*_A_⟩ = *p*_A_*K*.

We validate our predictions by simulating microbial community assembly, assuming *r*_A_ = 1 and *r*_B_ = 1.05, which corresponds to a selection coefficient of *s* = −0.05. Figure 3A shows the mean relative abundance of species A as a function of dispersal rate for various cluster sizes. In the high-dispersal regime, where community assembly is governed solely by dispersal, the mean relative abundance converges to that of the microbial pool, rendering replication rates insignificant. Similarly, in the low-dispersal regime, this convergence occurs when the cluster size is 1, as the first dispersing individual populates the entire community before a subsequent dispersal event, making replication rates inconsequential (Marrec and Bank, 2024). However, for cluster sizes greater than 1, dispersal introduces multiple species, leading to competitive dynamics when replication rates differ. As a result, the mean relative abundance, given by Equation 3, deviates from that of the microbial pool.

**Figure 3:**
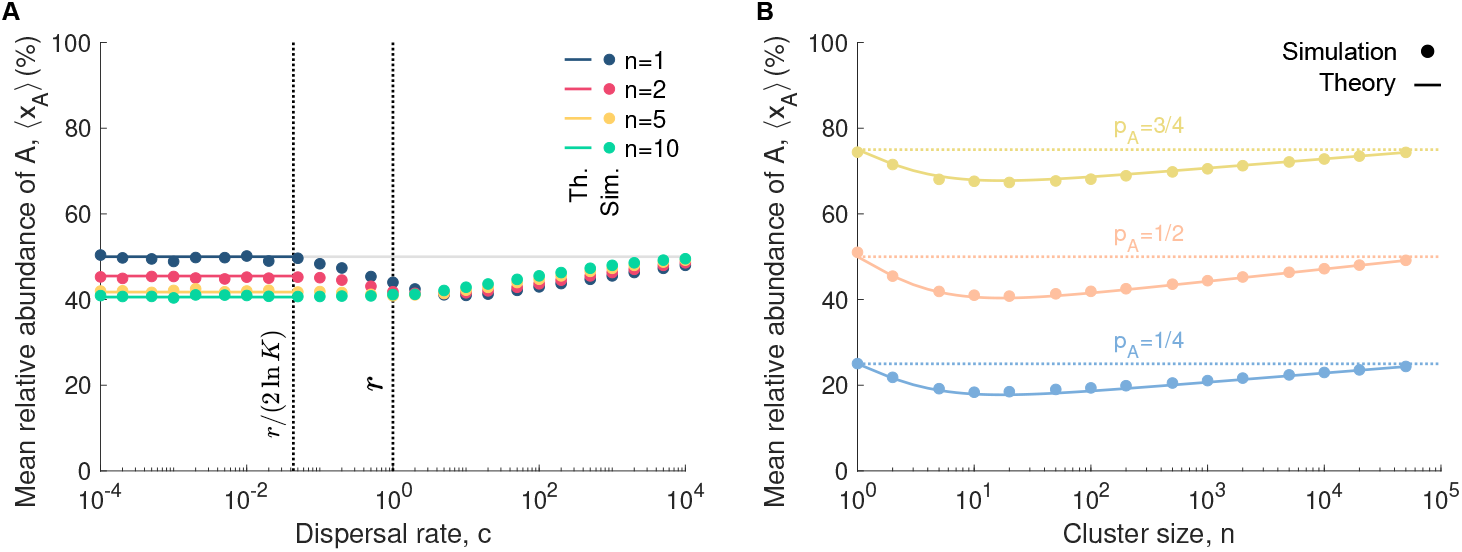
Cluster dispersal and within-community selection alter species abundance in a non-monotonic way. Panel **A** shows the mean relative abundance of A ⟨*x*_A_⟩ = ⟨*N*_A_⟩ */K* as a function of the dispersal rate *c* for various cluster sizes *n*, whereas panel **B** shows it as a function of the cluster size *n* for different abundances in the pool *p*_A_ in the low-dispersal regime. In both panels, the simulated data are averaged over 10^4^ microbial communities, whereas the solid lines represent our analytical predictions (Equations 3 and 4). Parameter values: replication rate of A *r*_A_ = 1, replication rate of B *r*_B_ = 1.05, dispersal rate *c* = 10^−4^ (in **B**), relative abundance of A in the pool *p*_A_ = 1*/*2 (in **A**), carrying capacity *K* = 10^5^.

Figure 3B depicts the mean relative abundance of species A as a function of cluster size in the low-dispersal regime, revealing the non-monotonic pattern described above, with an extremum at intermediate cluster sizes. For clusters of size 1, the probability of containing an A microbe is simply *p*_A_, and this single microbe populates the community without competition. Conversely, large clusters, which contain on average *p*_A_ × *n* A microbes dominate microbial community assembly due to their size, leaving limited opportunities for cell replication. In both extreme cases, replication rate differences become negligible. However, at intermediate cluster sizes, dispersal introduces a mix of species, leading to competition that amplifies the effects of differential replication rates, driving the observed non-monotony. These dynamics underscore the critical role of cluster dispersal in shaping community structure within the low-dispersal regime.

In summary, when cluster size equals one, selection influences community assembly only at intermediate dispersal rates. However, as soon as clusters contain more than one individual, selection also becomes relevant in the low-dispersal regime—unless the clusters are excessively large. Thus, cluster dispersal modulates the extent to which selection shapes microbial community assembly by influencing species abundance.

### Bimodality coefficient and species abundance shed light on assembly regime, within-community selection and cluster dispersal

Previous work demonstrated that the bimodality coefficient and mean relative abundance can serve as metrics to characterize microbial community assembly (Vega and Gore, 2017; Marrec and Bank, 2024). Specifically, the bimodality coefficient helps determine whether assembly is driven by cell replication, dispersal events, or both, while mean relative abundance reflects differences in microbial traits, such as replication rates.

Figures 4 and S2 present the mean relative abundance as a function of the bimodality coefficient for data simulated with various microbial pool abundances *p*_A_ and selection coefficients *s* = *r*_A_−*r*_B_. Regardless of the microbial pool or selection coefficient, the bimodality coefficient decreases to 1/3 as the dispersal rate increases. The mean relative abundance provides insights into the values of *p*_A_ and *s*. When selection is present, that is, when the selection coefficient satisfies *s* ≠ 0, the mean relative abundance of species A becomes a non-monotonic function of the bimodality coefficient and displays an extremum. Specifically, when species A is beneficial (*s >* 0), the curve exhibits a maximum, while when it is deleterious (*s <* 0), it shows a minimum. In contrast, under neutral conditions (*s* = 0), the relationship is flat. Moreover, if species A is more prevalent in the microbial pool, its mean relative abundance exceeds 50%. If it is less prevalent, the mean falls below 50%.

**Figure 4:**
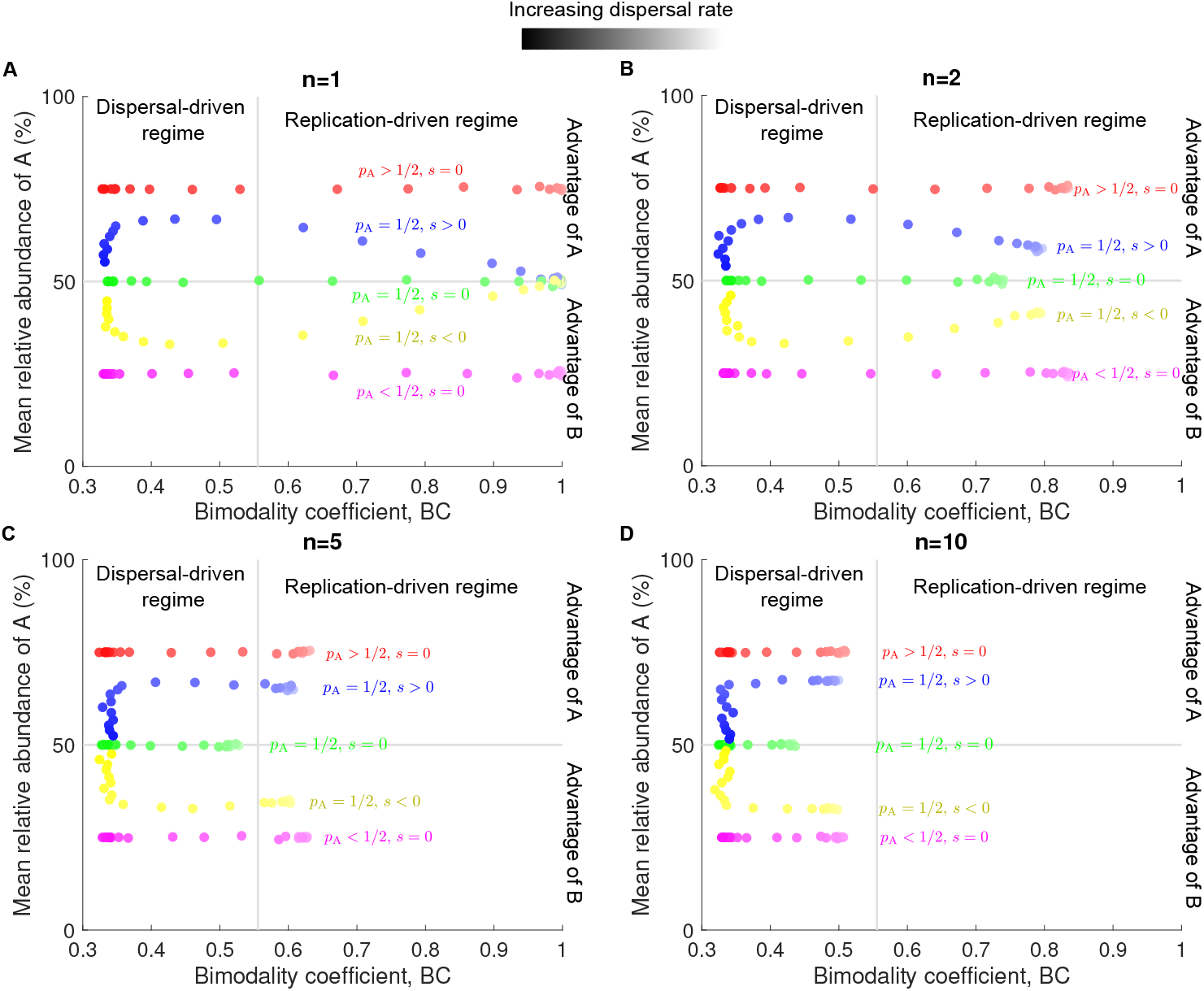
Bimodality coefficient and mean relative abundance reveal patterns of selection and cluster dispersal. Each panel shows the mean relative abundance of A as a function of the bimodality coefficient for different cluster sizes *n*. In all panels, the simulated data are averaged over 10^4^ stochastic replicates (i.e., microbial communities). Parameter values: replication rate of A *r*_A_ = 1 (red, green, yellow, purple) *r*_A_ = 1.1 (blue), replication rate of B *r*_B_ = 1 (red, blue, green, purple) *r*_B_ = 1.1 (yellow), relative abundance of A in the pool *p*_A_ = 1*/*2 (blue, green, yellow) *p*_A_ = 3*/*4 (red) *p*_A_ = 1*/*4 (purple), carrying capacity *K* = 10^5^, dispersal rate *c* = 10^−4^ − 10^4^.

Importantly, we show that both metrics, the bimodality coefficient and mean relative abundance, are influenced by the number of microbes dispersing simultaneously, i.e., cluster size. In particular, as cluster size increases, the maximum value of the bimodality coefficient decreases (Figure 2), whereas its effect on mean relative abundance is non-monotonic (Figure 3). Nonetheless, our results demonstrate that, across different dispersal rates, these two metrics still reveal patterns that distinguish whether two species differ in abundance within the microbial pool and exhibit varying replication rates (Figure 4). When cluster size is large, a higher number of data points is required to avoid misidentifying, for example, cases (*p*_A_ = 1*/*2, *s <* 0) and (*p*_A_ *<* 1*/*2, *s* = 0). Figures 4C and 4D illustrate a case where mean relative abundance remains relatively constant across a wide range of dispersal rates, even when the selection coefficient is nonzero, a pattern not observed for small cluster sizes (Figures 4A and 4B).

### Cluster dispersal is detected in experimental data

We applied our approach to three datasets collected by Vega and Gore (2017), who investigated the assembly of the gut microbiota in *C. elegans* by feeding them mixtures of bacteria. The first dataset examines the gut microbiota of AU37 worm strains fed a 50/50 mixture of fluorescently labeled *E. coli* strains (YFP and dsRed) (Figure 5A). The second dataset mirrors the first, differing only in the worm strain, which is glp-4 (Figure 5B). The third dataset explores the assembly of the gut microbiota in AU37 worms using a 50/50 mixture of two distinct bacterial species, *S. marcescens* and *E. aerogenes* (Figure 5C). By varying mixture densities, Vega and Gore (2017) quantified microbial community structures across a range of dispersal rates.

**Figure 5:**
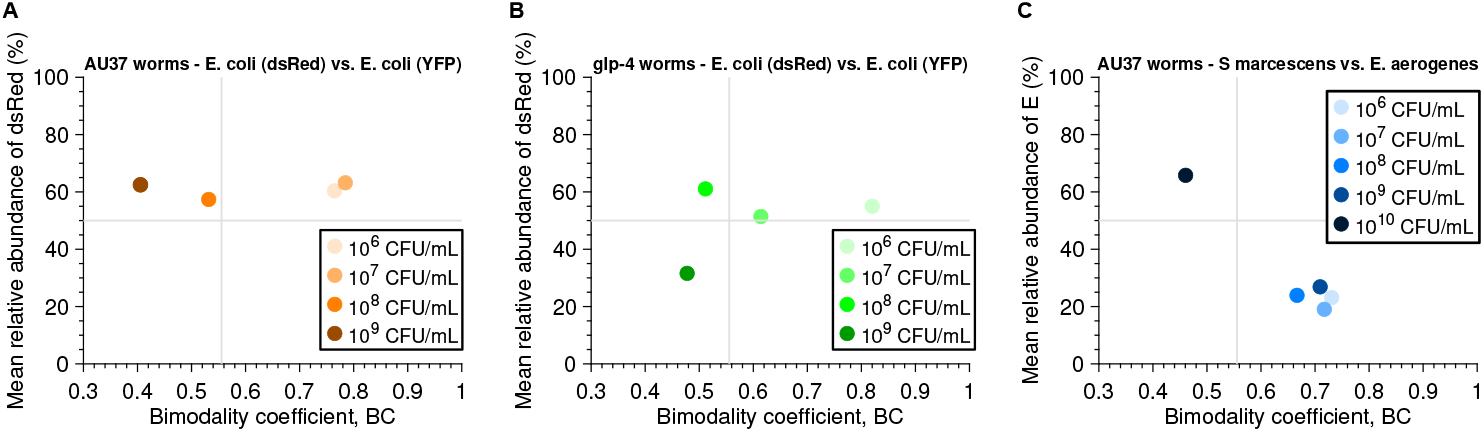
Bimodality coefficient and mean relative abundance reveal patterns of cluster dispersal in experimental data. Panels **A** and **B** show the mean relative abundance of *E. coli* (dsRed) as a function of the bimodality coefficient for different worm strains (AU37 in **A** and glp-4 in **B**). Panel **C** shows the mean relative abundance of *E*.*aerogenes* as a function of the bimodality coefficient for AU37 worms. Experimental data collected by Vega and Gore (2017).

Figure 5 shows the mean relative abundance as a function of the bimodality coefficient for their data sets. The first observation is that the bimodality coefficient decreases as the bacterial density of the environmental pool, and therefore the dispersal rate, increases. Interestingly, the maximum value of the bimodality coefficient obtained for these three data sets is around 0.8. From figures A and C, it is clear that several bacterial densities give the same bimodality coefficient, which corresponds to the plateau observed in the low-dispersal regime (Figure 2A). We can therefore assume that the cluster size is greater than 1.

The data presented in Figure 5A exhibit a pattern similar to that observed in Figure 4 for *p*_A_ *>* 1*/*2 and *s* = 0. This suggests that the dsRed strain is more abundant than the YFP strain in the microbial pool, while both strains appear to have similar replication rates. Using the equation ⟨*N*_A_⟩ = *p*_A_*K* and the relative abundance averaged over the four data points in Figure 5A, we estimate *p*_A_ ≈ 3*/*5. Applying Equation 3 in the low-dispersal regime and averaging the bimodality coefficient over the two data points corresponding to bacterial densities of 10^6^ and 10^7^ CFU/mL, we estimate *n* ≈ 2.

In Figure 5B, the relative abundance of three data points is close to 50%, except for the point corresponding to a bacterial density of 10^9^ CFU/mL. Based on this observation, we assume *p*_A_ = 1*/*2 and *s* = 0. Using Equation 3 in the low-dispersal regime and the bimodality coefficient obtained for a bacterial density of 10^6^ CFU/mL, we estimate *n* ≈ 1.6.

The data in Figure 5C are more challenging to interpret, as they do not follow any clear pattern observed in Figure 4. Notably, the relative abundance at a bacterial density of 10^6^ CFU/mL differs significantly from the other data points. Given this discrepancy, we exclude it from our analysis and focus on the remaining four data points. Assuming *s* = 0 and applying the same reasoning as for Figures 5A and B, we estimate *p*_A_ ≈ 1*/*5 and *n* ≈ 3.6.

Consequently, our predictions allowed us to shed light on evidence of cluster dispersal in experimental data and estimate cluster size.

### Our predictions remain valid with multiple species

So far, we have considered that the microbial pool contains only two species, which is relevant to some experimental data (Vega and Gore, 2017; Ortiz et al., 2021; Jones et al., 2022). However, in the wild, microbial communities are likely to be made up of multiple species. This leads us to extend our model to *S* species, assuming they all have the same replication rate *r*. These species are denoted by *i* = 1, 2, …, *S* and are each present in the microbial pool in abundance 1*/S*. Since the bimodality coefficient is not suitable for cases with more than two species, we use *α*- and *β*-diversity to quantify richness and between-community dissimilarity, respectively.

Here, *α*-diversity simply corresponds to the number of microbial species that are found in the microbial community when its size equals carrying capacity. In the high-dispersal regime, richness is expected to equal the number of species in the pool. In the low-dispersal regime, richness is equal to that of the first cluster dispersing from the pool. The probability that a cluster of size *n* has a richness of *α*_R_ is given by (Supporting Information - Section 6)

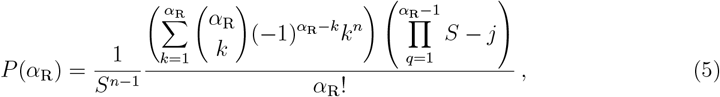

which allows us to derive the mean value of richness 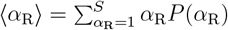.

To validate our analytical predictions, we generated *in silico* data by simulating the assembly of microbial communities from a pool containing seven neutral species. We then calculated their richness (Figure 6A and C). As shown in Figure 6A, richness increases with dispersal rate, regardless of cluster size. However, while high dispersal rates yield the same richness across cluster sizes, at low dispersal rates, larger clusters exhibit higher richness. In the low-dispersal regime, the richness of a microbial community reflects that of the first microbial cluster, whose composition may differ from the pool. Naturally, larger clusters may contain more species, leading to increased richness. In contrast, in the high-dispersal regime, microbial community composition mirrors that of the pool, resulting in uniform richness across cluster sizes.

**Figure 6:**
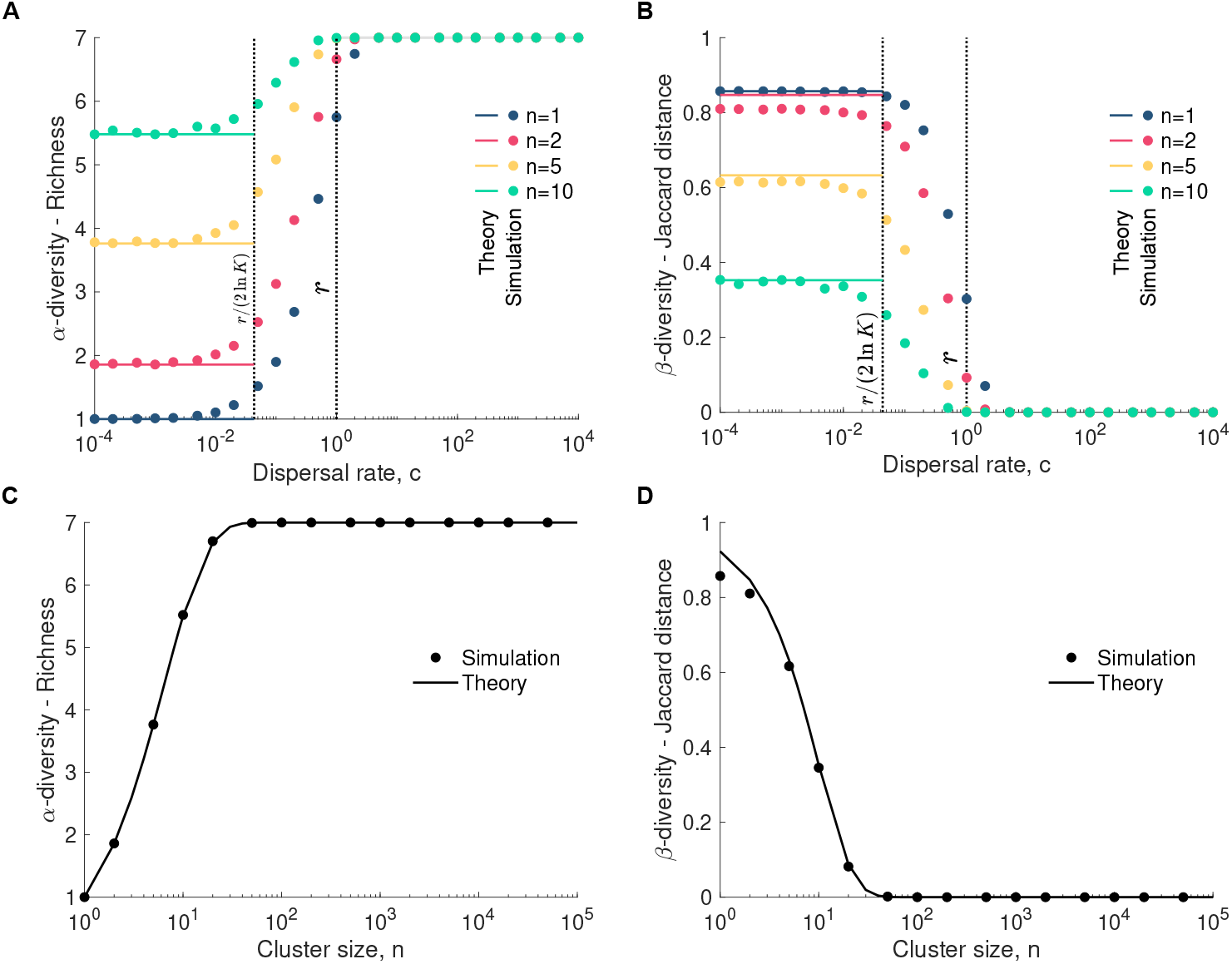
Cluster dispersal increases *α*-diversity and decreases *β*-diversity. Panels **A** and **C** show the richness as a function of the dispersal rate and cluster size, respectively, where **C** focuses on the low-dispersal regime. The simulated data are averaged over 10^3^ stochastic replicates (i.e., microbial communities). Panels **B** and **D** represent the Jaccard distance as a function of the dispersal rate and cluster size, respectively, where **D** focuses on the low-dispersal regime. The data points are averaged over the comparison of each pair of 10^3^ stochastic replicates (i.e., microbial communities). The solid lines represent our analytical predictions (Equations 5 and 8). Parameter values: replication rate *r* = 1, dispersal rate *c* = 10^−4^ (in **C** and **D**), number of species *S* = 7, relative abundance of each species in the pool *p* = 1*/S*, carrying capacity *K* = 10^5^.

In addition to *α*-diversity, we also quantify *β*-diversity, which determines the dissimilarity of two microbial communities. Here, we focus on the Jaccard distance, which quantifies the proportion of species that differ between two communities. Mathematically, the Jaccard distance is defined as *β*_J_ = 1 − *J* (X, Y), where *J* (X, Y) is the Jaccard similarity coefficient. This coefficient is equal to |X ∩ Y| */*(|X| + |Y| − |X ∩ Y|), where |X| and |Y| are the richness of communities X and Y, respectively, and |X ∩ Y| the number of shared species between communities X and Y. A Jaccard distance of 0 indicates identical composition, while a value of 1 signifies completely distinct communities.

In the high-dispersal regime, dispersal tends to homogenize community composition, resulting in a Jaccard distance close to zero. In contrast, in the low-dispersal regime, the Jaccard distance is influenced by the composition of the initial clusters that populate each community. Assuming that *p*_*i*_ = 1*/S* for 1 ≤ *i* ≤ *S*, we can show the probability that two communities have a number of shared species equal to |X ∩ Y| = *k* is approximately given by (Supporting Information - Section 6)

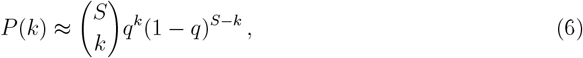

where

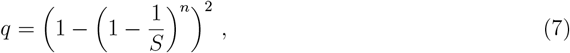

is the probability that a given species is present in both clusters and, thus, in both communities. Equation 6 allows us to compute the mean number of shared species 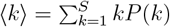. Thus, the mean Jaccard distance is approximately given by

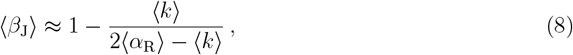

where we assumed that |X| and |Y| are simply equal to the mean richness ⟨*α*_R_⟩ .

Using the microbial communities generated previously, we compare each pair of them and compute their *β*-diversity. Figure 6 shows that *β*-diversity decreases with dispersal rate. In the high-dispersal regime, dispersal homogenizes microbial communities so they all have compositions similar to the microbial pool. In the low-dispersal regime, the larger cluster size, the lower *β*-diversity, as they likely introduce multiple species into microbial communities (Figure 6D).

## Discussion

In this work, we built a microbial community assembly model to examine the impact of cluster dispersal on community structure. Our model accounts for two events: dispersal and intra-community replication (Figure 1). The timescales associated with these two events strongly impact richness and dissimilarity during the stochastic assembly of microbial communities. In particular, Vega and Gore (2017) and Marrec and Bank (2024)’s work shed light on distinct assembly regimes: one driven by high dispersal rates relative to replication rates that homogenizes microbial communities, thus reducing between-community diversity while increasing richness. Conversely, low dispersal rates relative to replication rates are a barrier to high richness and lead to dissimilar structures. Here, we showed that cluster dispersal mitigates the latter effect. Specifically, clusters, when large, likely introduce multiple species into local communities at a time. Thus, even in the low-dispersal regime, dispersal can contribute to homogenizing microbial communities (Figure 2), showing that timescales are not the only parameter to consider when assessing the impact of dispersal on richness and between-community dissimilarity.

It is widely accepted that very low dispersal rates can lead to dispersal limitation, resulting in highly dissimilar microbial community structures (MacArthur and Wilson, 2001; Etienne and Olff, 2004; Zhou and Ning, 2017). Conversely, very high dispersal rates promote homogenization, leading to more similar community structures (MacArthur and Wilson, 2001; Etienne and Olff, 2004; Zhou and Ning, 2017). However, our study reveals that even at very low dispersal rates, microbial communities can become less dissimilar if microbes disperse in large clusters. This suggests that dispersal play an increasingly significant role as cluster size grows, regardless of dispersal rate, underscoring cluster size as a key factor in microbial community assembly and diversity.

We found evidence of cluster dispersal in experimental data collected by Vega and Gore (2017). They investigated the stochastic assembly of the gut microbiota in *Caenorhabditis elegans* worms by feeding them a microbial mixture. In a low-dispersal regime with two species, the expected bimodality coefficient is 1. However, we observed that Vega and Gore (2017)’s data yielded bimodality coefficient values plateauing around 0.8 (Figure 5). Our reanalysis, informed by our analytical predictions, revealed that in Vega and Gore (2017)’s experiment, worms likely ingested multiple microbes at once.

Marrec and Bank (2024) demonstrated how the bimodality coefficient and mean relative abundance can be used as metrics to identify assembly regimes and compare microbial traits across species in experimental data under various dispersal rates. Here, we extend their approach to scenarios beyond the dispersal of individual microbes (Figure 4). Specifically, while our results confirm that these two metrics consistently distinguish whether two species differ in abundance within the microbial pool or exhibit varying replication rates, a larger number of data points is required to prevent misidentification when cluster sizes are large.

Although multiple community assembly experiments involve two species (Vega and Gore, 2017; Ortiz et al., 2021; Jones et al., 2022), natural microbial communities likely harbor many more species. This led us to extend our model to *S* species. Since the bimodality coefficient is not suitable for cases with more than two species, we derived analytical predictions for *α*- and *β*-diversity, which allowed us to show that our results drawn for two species still hold for *S* species (Figure 6). Specifically, cluster dispersal homogenizes microbial communities increasing their richness and decreasing their between-community dissimilarity.

In summary, our work underscores the crucial role of cluster dispersal in the assembly of microbial communities, not only by influencing richness and between-community dissimilarity, but also by modulating the extent to which selection shapes community structure.

## Author Contributions

LM designed the study; LM performed the numerical and analytical work; LM and SL analyzed and interpreted the data; LM wrote the manuscript; LM and SL edited the manuscript.

## Conflict of interest declaration

The authors declare they have no competing interests.

## Funding

This research was supported as a part of NCCR Microbiomes, funded by the Swiss National Science Foundation (grant number 51NF40_ 225148).

## Acknowledgments

The authors thank the EE Group at Unil for insightful discussions. LM thanks Claudia Bank for discussions that led to the onset of the project.

## Notes

### Competing Interest Statement

The authors have declared no competing interest.

## References

A. Baud, K.-H. Hillion, C. Plainvert, V. Tessier, A. Tazi, L. Mandelbrot, C. Poyart, and S. P. Kennedy. Microbial diversity in the vaginal microbiota and its link to pregnancy outcomes. Scientific Reports, 13(1), June 2023. ISSN 2045-2322. doi: 10.1038/s41598-023-36126-z. URL http://dx.doi.org/10.1038/s41598-023-36126-z.

J. E. Blum, C. N. Fischer, J. Miles, and J. Handelsman. Frequent replenishment sustains the beneficial microbiome of drosophila melanogaster. mBio, 4(6), Dec. 2013. ISSN 2150-7511. doi: 10.1128/mbio.00860-13. URL http://dx.doi.org/10.1128/mbio.00860-13.

J. Clobert, E. Danchin, A. A. Dhondt, and J. D. Nichols, editors. Dispersal. Oxford University Press, London, England, Feb. 2001.

A. M. Ellison. Effect of seed dimorphism on the density-dependent dynamics of experimental populations of atriplex triangularis (chenopodiaceae). American Journal of Botany, 74(8): 1280–1288, Aug. 1987. ISSN 1537-2197. doi: 10.1002/j.1537-2197.1987.tb08741.x. URL http://dx.doi.org/10.1002/j.1537-2197.1987.tb08741.x.

R. S. Etienne and H. Olff. A novel genealogical approach to neutral biodiversity theory. Ecology Letters, 7(3):170–175, Feb. 2004. ISSN 1461-0248. doi: 10.1111/j.1461-0248.2004.00572.x. URL http://dx.doi.org/10.1111/j.1461-0248.2004.00572.x.

D. T. Gillespie. A general method for numerically simulating the stochastic time evolution of coupled chemical reactions. Journal of Computational Physics, 22(4):403–434, Dec. 1976. doi: 10.1016/0021-9991(76)90041-3. URL https://doi.org/10.1016/0021-9991(76)90041-3.

D. T. Gillespie. Exact stochastic simulation of coupled chemical reactions. The Journal of Physical Chemistry, 81(25):2340–2361, Dec. 1977. doi: 10.1021/j100540a008. URL https://doi.org/10.1021/j100540a008.

J. K. Goodrich, J. L. Waters, A. C. Poole, J. L. Sutter, O. Koren, R. Blekhman, M. Beaumont, W. Van Treuren, R. Knight, J. T. Bell, T. D. Spector, A. G. Clark, and R. E. Ley. Human genetics shape the gut microbiome. Cell, 159(4):789–799, Nov. 2014. ISSN 0092-8674. doi: 10.1016/j.cell.2014.09.053. URL http://dx.doi.org/10.1016/j.cell.2014.09.053.

J. Grilli. Macroecological laws describe variation and diversity in microbial communities. Nature Communications, 11(1), Sept. 2020. ISSN 2041-1723. doi: 10.1038/s41467-020-18529-y. URL http://dx.doi.org/10.1038/s41467-020-18529-y.

N. Hadland, C. W. Hamilton, and S. Duhamel. Young volcanic terrains are windows into early microbial colonization. Communications Earth and Environment, 5(1), Mar. 2024. ISSN 2662-4435. doi: 10.1038/s43247-024-01280-3. URL http://dx.doi.org/10.1038/s43247-024-01280-3.

B. Houchmandzadeh. Giant fluctuations in logistic growth of two species competing for limited resources. Physical Review E, 98(4), Oct. 2018. doi: 10.1103/physreve.98.042118. URL https://doi.org/10.1103/physreve.98.042118.

E. W. Jones, J. M. Carlson, D. A. Sivak, and W. B. Ludington. Stochastic microbiome assembly depends on context. Proceedings of the National Academy of Sciences, 119(7), Feb. 2022. doi: 10.1073/pnas.2115877119. URL https://doi.org/10.1073/pnas.2115877119.

O. F. A. Larsen and E. Claassen. The mechanistic link between health and gut microbiota diversity. Scientific Reports, 8(1), Feb. 2018. ISSN 2045-2322. doi: 10.1038/s41598-018-20141-6. URL http://dx.doi.org/10.1038/s41598-018-20141-6.

D. S. Lundberg, S. L. Lebeis, S. H. Paredes, S. Yourstone, J. Gehring, S. Malfatti, J. Tremblay, A. Engelbrektson, V. Kunin, T. G. d. Rio, R. C. Edgar, T. Eickhorst, R. E. Ley, P. Hugenholtz, S. G. Tringe, and J. L. Dangl. Defining the core arabidopsis thaliana root microbiome. Nature, 488(7409):86–90, Aug. 2012. ISSN 1476-4687. doi: 10.1038/nature11237. URL http://dx.doi.org/10.1038/nature11237.

R. MacArthur and E. Wilson. The Theory of Island Biogeography. Book collections on Project MUSE. Princeton University Press, 2001. ISBN 9780691088365. URL https://books.google.ch/books?id=a10cdkywhVgC.

L. Marrec and C. Bank. Drivers of diversity within and between microbial communities during stochastic assembly. Nov. 2024. doi: 10.1101/2024.11.19.624346. URL http://dx.doi.org/10.1101/2024.11.19.624346.

B. Obadia, Z. Güvener, V. Zhang, J. A. Ceja-Navarro, E. L. Brodie, W. W. Ja, and W. B. Ludington. Probabilistic invasion underlies natural gut microbiome stability. Current Biology, 27(13):1999–2006.e8, July 2017. ISSN 0960-9822. doi: 10.1016/j.cub.2017.05.034. URL http://dx.doi.org/10.1016/j.cub.2017.05.034.

A. Ortiz, N. M. Vega, C. Ratzke, and J. Gore. Interspecies bacterial competition regulates community assembly in the c. elegans intestine. The ISME Journal, 15(7):2131–2145, Feb. 2021. doi: 10.1038/s41396-021-00910-4. URL https://doi.org/10.1038/s41396-021-00910-4.

C. C. R. Smith, L. K. Snowberg, J. Gregory Caporaso, R. Knight, and D. I. Bolnick. Dietary input of microbes and host genetic variation shape among-population differences in stickle-back gut microbiota. The ISME Journal, 9(11):2515–2526, Apr. 2015. ISSN 1751-7370. doi: 10.1038/ismej.2015.64. URL http://dx.doi.org/10.1038/ismej.2015.64.

A. Spor, O. Koren, and R. Ley. Unravelling the effects of the environment and host genotype on the gut microbiome. Nature Reviews Microbiology, 9(4):279–290, Mar. 2011. ISSN 1740-1534. doi: 10.1038/nrmicro2540. URL http://dx.doi.org/10.1038/nrmicro2540.

K. Tillisch, J. Labus, L. Kilpatrick, Z. Jiang, J. Stains, B. Ebrat, D. Guyonnet, S. Legrain–Raspaud, B. Trotin, B. Naliboff, and E. A. Mayer. Consumption of fermented milk product with probiotic modulates brain activity. Gastroenterology, 144(7): 1394–1401.e4, June 2013. ISSN 0016-5085. doi: 10.1053/j.gastro.2013.02.043. URL http://dx.doi.org/10.1053/j.gastro.2013.02.043.

A. Tsoularis and J. Wallace. Analysis of logistic growth models. Mathematical Biosciences, 179(1):21–55, July 2002. doi: 10.1016/s0025-5564(02)00096-2. URL https://doi.org/10.1016/s0025-5564(02)00096-2.

E. van Nood, A. Vrieze, M. Nieuwdorp, S. Fuentes, E. G. Zoetendal, W. M. de Vos, C. E. Visser, E. J. Kuijper, J. F. Bartelsman, J. G. Tijssen, P. Speelman, M. G. Dijkgraaf, and J. J. Keller. Duodenal infusion of donor feces for recurrentclostridium difficile. New England Journal of Medicine, 368(5):407–415, Jan. 2013. ISSN 1533-4406. doi: 10.1056/nejmoa1205037. URL http://dx.doi.org/10.1056/NEJMoa1205037.

N. M. Vega and J. Gore. Stochastic assembly produces heterogeneous communities in the caenorhabditis elegans intestine. PLOS Biology, 15(3):e2000633, Mar. 2017. doi: 10.1371/journal.pbio.2000633. URL https://doi.org/10.1371/journal.pbio.2000633.

R. Vilchez-Vargas, J. Skieceviciene, K. Lehr, G. Varkalaite, C. Thon, M. Urba, E. Morkunas, L. Kucinskas, K. Bauraite, D. Schanze, M. Zenker, P. Malfertheiner, J. Kupcinskas, and A. Link. Gut microbial similarity in twins is driven by shared environment and aging. eBioMedicine, 79:104011, May 2022. ISSN 2352-3964. doi: 10.1016/j.ebiom.2022.104011. URL http://dx.doi.org/10.1016/j.ebiom.2022.104011.

R. Zapién-Campos, M. Sieber, and A. Traulsen. Stochastic colonization of hosts with a finite lifespan can drive individual host microbes out of equilibrium. PLOS Computational Biology, 16(11):e1008392, Nov. 2020. doi: 10.1371/journal.pcbi.1008392. URL https://doi.org/10.1371/journal.pcbi.1008392.

F. Zhang, M. Berg, K. Dierking, M.-A. Félix, M. Shapira, B. S. Samuel, and H. Schulenburg. Caenorhabditis elegans as a model for microbiome research. Frontiers in Microbiology, 8, Mar. 2017. ISSN 1664-302X. doi: 10.3389/fmicb.2017.00485. URL http://dx.doi.org/10.3389/fmicb.2017.00485.

J. Zhou and D. Ning. Stochastic community assembly: Does it matter in microbial ecology? Microbiology and Molecular Biology Reviews, 81(4), Dec. 2017. ISSN 1098-5557. doi: 10.1128/mmbr.00002-17. URL http://dx.doi.org/10.1128/mmbr.00002-17.

