## Supporting Information for "Cluster dispersal shapes microbial diversity during community assembly"

**2 Abundance fluctuation distributions** **5**

**3 Effective dispersal rate** **6**

**4 Bimodality coefficient and selection** **7**

**5 Data analysis through bimodality coefficient and species abundance** **8**

**6 Extension to $S$ species** **9**

### 1 Formal analysis

The assembly of a microbial community is a stochastic process that involves, in our model, two events: dispersal of a cluster from the microbial pool to the community, and intra-community cell replication. As the community is initially microbe-free, its assembly starts with small sizes. A deterministic formalism, based on a system of ordinary differential equations, does not adequately capture the stochastic dynamics of community growth involving low sizes (Marrec et al., 2023).

To address this, we adopt a microscopic, probabilistic description of microbial community assembly. More specifically, we formulate a system of equations that governs the dynamics of the probability of having $N_\mathrm{A}$ microbes of strain A in a community of size $N$ (and thus $N_\mathrm{B} = N - N_\mathrm{A}$ microbes of strain B), denoted by $P(N_\mathrm{A}, N)$.

We distinguish between two regimes: the low- and high-dispersal regimes.

#### 1.1 Low-dispersal regime

In the low-dispersal regime, a first cluster of size $n$ disperses and assembly is then completed by cell replication. This regime leads to the following system of equations (Houchmandzadeh, 2018)

$$P(N_\mathrm{A}, N+1) = \alpha_N^{N_\mathrm{A}-1} P(N_\mathrm{A}-1, N) + (1-\alpha_N^{N_\mathrm{A}}) P(N_\mathrm{A}, N)\,, \tag{1}$$

with $N_{\mathrm{A},0}$ as an initial condition, which is drawn from the binomial dsitribution $\mathcal{B}(n, p_\mathrm{A})$. The function $\alpha_N^{N_\mathrm{A}}$ denotes the probability that the increase in community size by one microbe ($N \to N+1$) is due to the replication of an A microbe. This probability is given by (Houchmandzadeh, 2018)

$$\alpha_N^{N_\mathrm{A}} = \frac{r_\mathrm{A} N_\mathrm{A}}{r_\mathrm{A} N_\mathrm{A} + r_\mathrm{B}(N - N_\mathrm{A})}\,. \tag{2}$$

We are interested in deriving the moments of $N_\mathrm{A}$, i.e., $\langle N_\mathrm{A}^k \rangle = \sum_{N_\mathrm{A}=0}^{N} N_\mathrm{A}^k P(N_\mathrm{A}, N)$. They can be calculated by multiplying Equation 1 by $N_\mathrm{A}^k$, summing over the possible values of $N_\mathrm{A}$, and solving the resulting recurrence relation . Assuming the neutral case ($r_\mathrm{A} = r_\mathrm{B} = r$) simplifies the derivation of the moments, which leads to

$$\langle N_\mathrm{A} \rangle = p_\mathrm{A} N\,, \tag{3}$$

$$\langle N_\mathrm{A}^2 \rangle \underset{N\gg1}{\approx} \frac{(2+(n-1)p_\mathrm{A})p_\mathrm{A}N^2}{n+1}\,, \tag{4}$$

$$\langle N_\mathrm{A}^3 \rangle \underset{N\gg1}{\approx} \frac{(6+(n-1)p_\mathrm{A}(6+(n-2)p_\mathrm{A}))p_\mathrm{A}N^3}{(n+1)(n+2)}\,, \tag{5}$$

and

$$\langle N_\mathrm{A}^4 \rangle \underset{N\gg1}{\approx} \frac{(24+(n-1)p_\mathrm{A}(36+(n-2)p_\mathrm{A}(12+(n-3)p_\mathrm{A})))p_\mathrm{A}N^4}{(n+1)(n+2)(n+3)}\,. \tag{6}$$

Note that these moments were derived by first calculating the moments of $N_{\mathrm{A},0}$ (e.g., $\langle N_{\mathrm{A},0} \rangle = p_\mathrm{A} n$). Now we can derive the variance, skewness, and kurtosis, which are given by

$$\sigma^2 \underset{N\gg1}{\approx} \frac{2(1-p_\mathrm{A})p_\mathrm{A}N^2}{n+1}\,, \tag{7}$$

$$\gamma \underset{N\gg1}{\approx} \frac{3(1-2p_\mathrm{A})}{(n+2)\sqrt{\frac{2(1-p_\mathrm{A})p_\mathrm{A}}{1+n}}}\,, \tag{8}$$

and

$$\kappa \underset{N\gg 1}{\approx} \frac{(n+1)\left(p_\mathrm{A}\left(n^2 p_\mathrm{A}(p_\mathrm{A}(3p_\mathrm{A}-1)-1)+2n(p_\mathrm{A}-1)(5p_\mathrm{A}-7)-6(13(p_\mathrm{A}-2)p_\mathrm{A}+17)\right)+24\right)}{4(n+2)(n+3)(1-p_\mathrm{A})^2 p_\mathrm{A}}\,. \tag{9}$$

The skewness and kurtosis allow us to obtain the bimodality coefficient (Ellison, 1987; Vega and Gore, 2017)

$$\mathrm{BC} = \frac{\gamma^2+1}{\kappa}\,. \tag{10}$$

Although we have an explicit formula for the bimodality coefficient, it is too complex to be written here.

If both species have distinct replication rates ($r_\mathrm{A} \neq r_\mathrm{B}$), resulting in a nonzero selection coefficient ($s \neq 0$), the moments are more difficult to obtain. To get around this difficulty, we approximate Equation 1 by assuming that $N_\mathrm{A}$ and $N$ are continuous variables. This assumption allows us to reduce Equation 1 to the partial differential equation (Houchmandzadeh, 2018)

$$\frac{\partial P(N_\mathrm{A},N)}{\partial N} + \frac{\partial}{\partial N_\mathrm{A}}\left(\alpha_N^{N_\mathrm{A}} P(N_\mathrm{A},N)\right) = 0\,. \tag{11}$$

The moments $\langle N_\mathrm{A}^k\rangle$ can be derived from the previous equation. The neutral case ($r_\mathrm{A} = r_\mathrm{B}$) leads to (Houchmandzadeh, 2018)

$$\langle N_\mathrm{A}^k\rangle_\emptyset = \int_0^N N_\mathrm{A}^k P(N_\mathrm{A},N)\mathrm{d}N_\mathrm{A} = \frac{(N_{\mathrm{A},0})_k}{(n)_k}N^k\,. \tag{12}$$

where $\emptyset$ indicates the neutral case, $N_{\mathrm{A},0}$ denotes the number of A microbes in the first cluster, and $(x)_p = x(x+1)...(x+p-1)$ designates the Pochhammer symbol. To obtain the moments for nonzero selection coefficients, we apply a perturbation method, which, to the first order in $s$ ($s \ll 1$), yields

$$\langle N_\mathrm{A}^k\rangle = \sum_{N_{\mathrm{A},0}=0}^{n} \binom{n}{N_{\mathrm{A},0}} p_\mathrm{A}^{N_{\mathrm{A},0}}(1-p_\mathrm{A})^{n-N_{\mathrm{A},0}}\langle N_\mathrm{A}^k\rangle_\emptyset \left[1 + k\frac{n-N_{\mathrm{A},0}}{n+k}\log\left(\frac{N}{n}\right)s\right]. \tag{13}$$

As done previously, we can obtain the variance, skewness, and kurtosis from the moments, and then the bimodality coefficient.

#### 1.2 High-dispersal regime

In the high-dispersal regime, microbial community assembly is achieved only by dispersal events, without any cell replication. In this regime, a community reaches a size $N$ through the dispersal of $q$ clusters of size $n$, where $q = N/n$. Thus, the number of A microbes in the final structure of a community is equal to the sum of $q$ random variables that are independently drawn from the binomial distribution $\mathcal{B}(n, p_\mathrm{A})$. The sum of $q$ random variables that are drawn from the binomial distribution $\mathcal{B}(n, p_\mathrm{A})$ is itself drawn from the binomial distribution $\mathcal{B}(q \times n, p_\mathrm{A})$. Thus, the probability of having $N_\mathrm{A}$ A microbes in a community of size $N$ satisfies

$$P(N_\mathrm{A},N) = \binom{N}{N_\mathrm{A}} p_\mathrm{A}^{N_\mathrm{A}}(1-p_\mathrm{A})^{N-N_\mathrm{A}}\,, \tag{14}$$

where $N$ must be a multiple of $n$, otherwise the probability is zero. The first four moments satisfy

$$\langle N_\mathrm{A}\rangle = p_\mathrm{A} N\,, \tag{15}$$

$$\langle N_{\mathrm{A}}^2\rangle = p_{\mathrm{A}}(N + (N-1)Np_{\mathrm{A}})\,, \tag{16}$$

$$\langle N_{\mathrm{A}}^3\rangle = Np_{\mathrm{A}}((N-1)p_{\mathrm{A}}((N-2)p_{\mathrm{A}}+3)+1)\,, \tag{17}$$

and

$$\langle N_{\mathrm{A}}^4\rangle = Np_{\mathrm{A}}((N-1)p_{\mathrm{A}}((N-2)p_{\mathrm{A}}((N-3)p_{\mathrm{A}}+6)+7)+1)\,. \tag{18}$$

From the first four moments, we can derive the variance, skewness, and kurtosis, which are respectively given by

$$\sigma^2 = (1-p_{\mathrm{A}})p_{\mathrm{A}}N\,, \tag{19}$$

$$\gamma = \frac{1-2p_{\mathrm{A}}}{\sqrt{(1-p_{\mathrm{A}})p_{\mathrm{A}}N}}\,, \tag{20}$$

and

$$\kappa = 3 - \frac{6}{N} + \frac{1}{(1-p_{\mathrm{A}})p_{\mathrm{A}}N}\,. \tag{21}$$

The previous quantities allow us to derive an equation for the bimodality coefficient

$$\mathrm{BC} = \frac{1+(N-4)(1-p_{\mathrm{A}})p_{\mathrm{A}}}{1+3(N-2)(1-p_{\mathrm{A}})p_{\mathrm{A}}} \underset{N\gg 1}{\approx} \frac{1}{3}\,. \tag{22}$$

#### 1.3 Both regimes in the neutral case for a cluster size of 1

There is a special case that does not require both assembly regimes to be considered separately: the neutral case ($r_{\mathrm{A}} = r_{\mathrm{B}}$) with a cluster size of 1 ($n = 1$). In this case, the probability $\alpha_N^{N_{\mathrm{A}}}$, given by Equation 2, becomes

$$\alpha_N^{N_{\mathrm{A}}} = \frac{rN_{\mathrm{A}} + cp_{\mathrm{A}}}{rN + c}\,, \tag{23}$$

while the initial condition of the probability $P(N_{\mathrm{A}}, N)$, given by Equation 1, satisfies $P(0,0) = 1$. Here again, we can derive the first four moments of $N_{\mathrm{A}}$. They are given by

$$\langle N_{\mathrm{A}}\rangle = p_{\mathrm{A}}N\,, \tag{24}$$

$$\langle N_{\mathrm{A}}^2\rangle \underset{N\gg 1}{\approx} \frac{(cp_{\mathrm{A}}+r)p_{\mathrm{A}}N^2}{c+r}\,, \tag{25}$$

$$\langle N_{\mathrm{A}}^3\rangle \underset{N\gg 1}{\approx} \frac{(cp_{\mathrm{A}}+r)(cp_{\mathrm{A}}+2r)p_{\mathrm{A}}N^3}{(c+r)(c+2r)}\,, \tag{26}$$

and

$$\langle N_{\mathrm{A}}^4\rangle \underset{N\gg 1}{\approx} \frac{(cp_{\mathrm{A}}+r)(cp_{\mathrm{A}}+2r)(cp_{\mathrm{A}}+3r)p_{\mathrm{A}}N^4}{(c+r)(c+2r)(c+3r)}\,. \tag{27}$$

From the previous moments, we can derive the variance, skewness, and kurtosis, which are respectively given by

$$\sigma^2 \underset{N\gg 1}{\approx} \frac{r(1-p_{\mathrm{A}})p_{\mathrm{A}}N^2}{c+r}\,, \tag{28}$$

$$\gamma \underset{N\gg 1}{\approx} \frac{2r(1-2p_{\mathrm{A}})}{(c+2r)\sqrt{\frac{r(1-p_{\mathrm{A}})p_{\mathrm{A}}}{c+r}}}\,, \tag{29}$$

and

$$\kappa \underset{N\gg 1}{\approx} \frac{3(c+r)(2r+(c-6r)(1-p_{\mathrm{A}})p_{\mathrm{A}})}{(c+2r)(c+3r)(1-p_{\mathrm{A}})p_{\mathrm{A}}}\,. \tag{30}$$

Finally, we derive the bimodality coefficient

$$\mathrm{BC} \underset{N\gg 1}{\approx} \frac{(c+3r)\left(c^2(p_{\mathrm{A}}-1)p_{\mathrm{A}} - 4c(3(p_{\mathrm{A}}-1)p_{\mathrm{A}}+1)r - 4(3(p_{\mathrm{A}}-1)p_{\mathrm{A}}+1)r^2\right)}{3(c+r)(c+2r)((p_{\mathrm{A}}-1)p_{\mathrm{A}}(c-6r)-2r)}\,. \tag{31}$$

### 2 Abundance fluctuation distributions

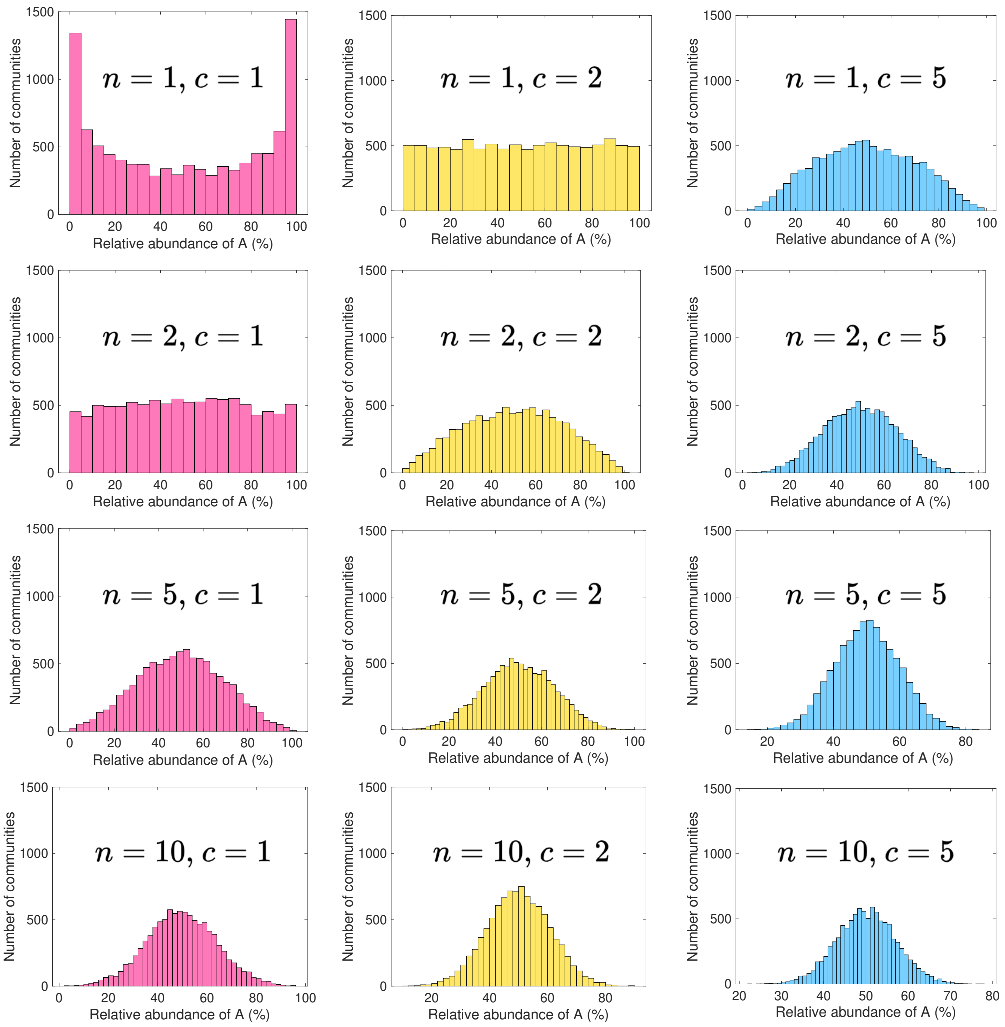

Figure 1: **Cluster dispersal homogenizes microbial communities.** Each panel shows the number of communities as a function of the relative abundance of A for a given pair $(n,\ c)$, where $n$ is the cluster size and $c$ is the dispersal rate. Parameter values : replication rates $r_{\mathrm{A}} = r_{\mathrm{B}} = 1$, carrying capacity $K = 10^5$, abundance of species A in the pool $p_{\mathrm{A}} = 1/2$, number of communities $10^3$.

### 3 Effective dispersal rate

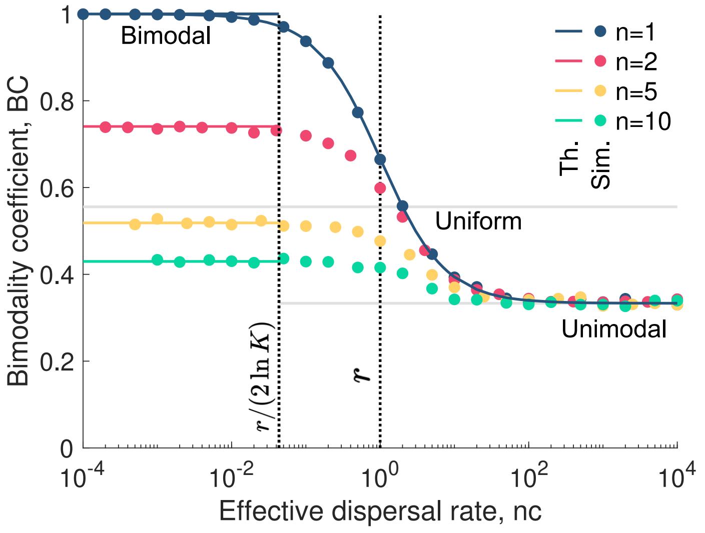

Figure 2: **Considering cluster dispersal is not equivalent to increasing the dispersal rate of individual microbes.** Bimodality coefficient BC as a function of the effective dispersal rate $n \times c$ for various cluster sizes $n$. The effective dispersal rate is the dispersal of clusters $c$ times their size $n$. The simulated data are averaged over $10^4$ microbial communities. The solid lines represent our analytical predictions (see main text). Parameter values: replication rates $r_\mathrm{A} = r_\mathrm{B} = 1$, relative abundance of A in the pool $p_\mathrm{A} = 1/2$, carrying capacity $K = 10^5$.

### 4 Bimodality coefficient and selection

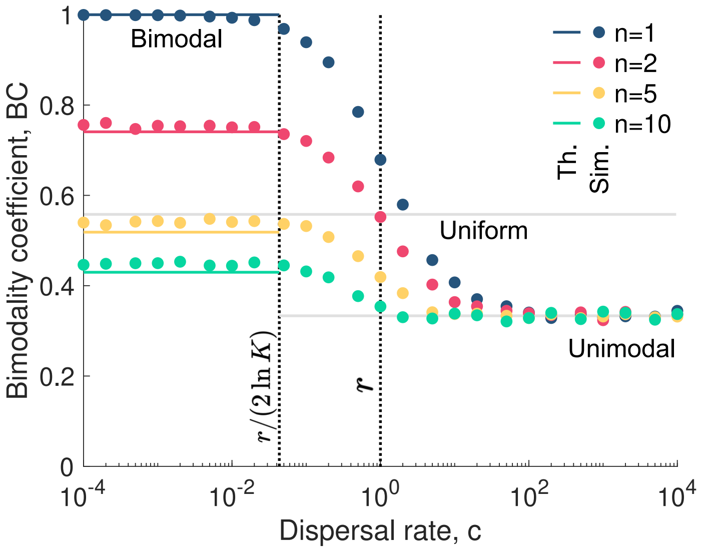

Figure 3: **Selection induces bimodality coefficient values similar to those observed under neutrality.** Bimodality coefficient BC as a function of the dispersal rate $c$ for various cluster sizes $n$. The simulated data are averaged over $10^4$ microbial communities. The solid lines represent our analytical predictions (see main text). Parameter values: replication rate of A $r_{\mathrm{A}} = 1$, replication rate of B $r_{\mathrm{B}} = 1.05$ relative abundance of A in the pool $p_{\mathrm{A}} = 1/2$, carrying capacity $K = 10^5$.

### 5 Data analysis through bimodality coefficient and species abundance

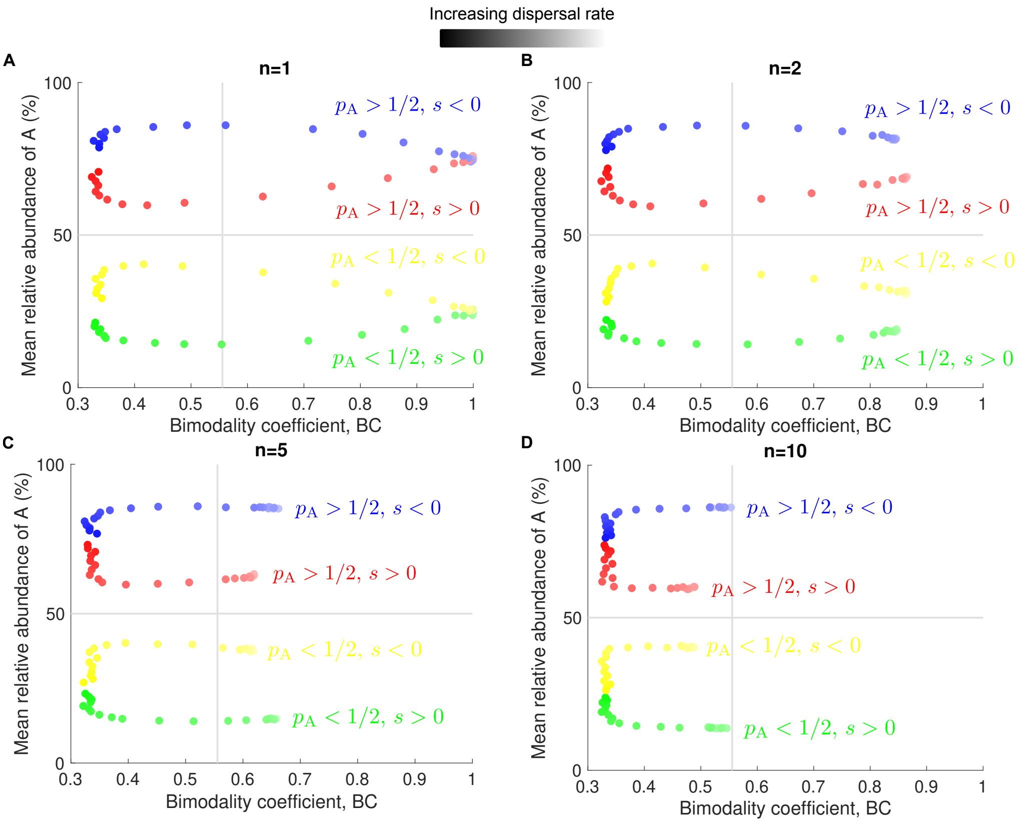

Figure 4: **Bimodality coefficient and mean relative abundance reveal patterns of cluster dispersal.** Each panel shows the mean relative abundance of A as a function of the bimodality coefficient for different cluster sizes $n$. In all panels, the simulated data are averaged over $10^4$ stochastic replicates. Parameter values: replication rate of A $r_A = 1$ (blue, yellow) $r_A = 1.1$ (red, green), replication rate of B $r_B = 1$ (red, green) $r_B = 1.1$ (blue, yellow), relative abundance of A in the pool $p_A = 3/4$ (blue, red) $p_A = 1/4$ (yellow, green), carrying capacity $K = 10^5$, dispersal rate $c = 10^{-4} - 10^4$.

### 6 Extension to $S$ species

In the main text, we consider a microbial pool composed of two species. Here, we generalize the model to include $S$ species, all assumed to replicate at the same rate $r$. These species are indexed by $i = 1, 2, \ldots, S$ and are present in the microbial pool with respective abundances $p_1, p_2, \ldots, p_S$.

Since the bimodality coefficient is not applicable for microbial communities with more than two species, we quantify diversity using two measures: $\alpha$-diversity, which captures within-community richness, and $\beta$-diversity, which describes between-community dissimilarity.

#### 6.1 $\alpha$-diversity

##### 6.1.1 Richness

In this work, we define $\alpha$-diversity as the number of distinct microbial species present in a community when it reaches carrying capacity, i.e., richness. We denote this quantity by $\alpha_{\mathrm{R}}$.

##### 6.1.2 High-dispersal regime

In the limit of high dispersal, the richness of each community is expected to match the full species pool, i.e., $\alpha_{\mathrm{R}} = S$.

##### 6.1.3 Low-dispersal regime

In the low-dispersal regime, community richness corresponds to the number of species present in the initial cluster that disperses from the pool.

**Multinomial sampling.** The species composition of a cluster of size $n$ is drawn from a multinomial distribution given by

$$P(N_1, N_2, \ldots, N_S) = \frac{n!}{N_1! N_2! \cdots N_S!} p_1^{N_1} p_2^{N_2} \cdots p_S^{N_S}, \tag{32}$$

where $N_i$ is the number of microbes of species $i$ in the cluster, subject to the constraint

$$\sum_{i=1}^{S} N_i = n, \tag{33}$$

ensuring the total number of microbes equals the cluster size. Additionally, the abundances in the pool satisfy

$$\sum_{i=1}^{S} p_i = 1. \tag{34}$$

**Clusters with a single species.** Consider a cluster composed solely of species $i$, i.e., all $n$ microbes belong to species $i$. The probability of such a cluster is

$$P(N_1 = 0, \ldots, N_i = n, \ldots, N_S = 0) = p_i^n. \tag{35}$$

Summing over all $S$ species, the probability that a cluster contains only one species is

$$P(\alpha_{\mathrm{R}} = 1) = \sum_{i=1}^{S} p_i^n. \tag{36}$$

**Clusters with two species.** Next, consider a cluster containing exactly two species, $i$ and $j$. Such a cluster contains $N_i$ individuals of species $i$ and $n-N_i$ of species $j$, where $1 \le N_i \le n-1$. The probability for such a cluster is

$$\sum_{k=1}^{n-1} P(N_1 = 0, ..., N_i, ..., N_j = n - N_i, ..., N_S = 0) = \sum_{k=1}^{n-1} \frac{n!}{N_i!(n-N_i)!} p_i^{N_i} p_j^{n-N_i} = (p_i+p_j)^n - p_i^n - p_j^n \,. \tag{37}$$

Summing over all distinct pairs of species, the probability that a cluster contains exactly two species is

$$P(\alpha_{\rm R} = 2) = \sum_{i=1}^{S-1} \sum_{j=i+1}^{S} \left[ (p_i + p_j)^n - p_i^n - p_j^n \right] . \tag{38}$$

Assuming a uniform pool with $p_i = 1/S$ for all $i$, this expression simplifies to

$$P(\alpha_{\rm R} = 2) = \frac{1}{2}(2^n - 2)(S-1)\left(\frac{1}{S}\right)^{n-1} . \tag{39}$$

**General Case.** Following the reasoning above, one can derive a general expression for the probability $P(\alpha_{\rm R})$ of observing any particular richness level $\alpha_{\rm R} \in \{1, 2, \ldots, n\}$ (see Equation 5 in the main text).

#### 6.2 $\beta$-diversity

In addition to quantifying $\alpha$-diversity, we also assess $\beta$-diversity, which measures the dissimilarity between a pair of microbial communities.

##### 6.2.1 Jaccard distance

Here, we focus on the Jaccard distance, a metric that captures the proportion of species that differ between two communities. Mathematically, the Jaccard distance is defined as

$$\beta_{\rm J} = 1 - J({\rm X}, {\rm Y}) \,, \tag{40}$$

where $J({\rm X},{\rm Y}) = |{\rm X} \cap {\rm Y}|/|{\rm X} \bigcup {\rm Y}|$ is the Jaccard similarity coefficient. This coefficient represents the ratio of the number of shared species between communities X and Y to the total number of unique species across both communities. Equivalently, it can be expressed as

$$J({\rm X}, {\rm Y}) = |{\rm X} \cap {\rm Y}|/(|{\rm X}| + |{\rm Y}| - |{\rm X} \cap {\rm Y}|) \,, \tag{41}$$

where |X| and |Y| denote the species richness of communities X and Y, respectively. A Jaccard distance of 0 indicates identical species composition, while a value of 1 signifies completely distinct communities.

##### 6.2.2 High-dispersal regime

In the high-dispersal regime, dispersal tends to homogenize community composition, resulting in a Jaccard distance close to zero.

##### 6.2.3 Low-dispersal regime

In contrast, in the low-dispersal regime, the Jaccard distance is influenced by the composition of the initial clusters that populate each community. This case can be modeled by comparing the compositions of two clusters independently sampled from the same multinomial distribution

$$(N_{\mathrm{X},1}, N_{\mathrm{X},2}, ..., N_{\mathrm{X},S}) \sim \mathrm{Multinomial}(n, [p_1, p_2, ..., p_S]) \,, \tag{42}$$

and

$$(N_{\mathrm{Y},1}, N_{\mathrm{Y},2}, ..., N_{\mathrm{Y},S}) \sim \mathrm{Multinomial}(n, [p_1, p_2, ..., p_S]) \,, \tag{43}$$

where $N_{\mathrm{X},1}$ is the number of individuals of species 1 in the first cluster that disperses into community X, and similarly for the other variables.

For a given species $i$, the probability that both clusters contain at least one microbe of that species is

$$q_i = P(N_{\mathrm{X},i} > 0 \;\&\; N_{\mathrm{Y},i} > 0) = (1 - (1 - p_i)^n)^2 \,. \tag{44}$$

Assuming uniform species abundances, i.e., $p_i = 1/S$ for all $i$, the number of shared species $|\mathrm{X} \cap \mathrm{Y}| = k$ can be approximated by the following binomial distribution

$$P(k) \approx \binom{S}{k} q^k (1 - q)^{S-k} \,, \tag{45}$$

where $q = (1 - (1 - 1/S)^n)^2$ is the probability that a given species is present in both clusters.

### References

A. M. Ellison. Effect of seed dimorphism on the density-dependent dynamics of experimental populations of atriplex triangularis (chenopodiaceae). *American Journal of Botany*, 74(8): 1280–1288, Aug. 1987. ISSN 1537-2197. doi: 10.1002/j.1537-2197.1987.tb08741.x. URL http://dx.doi.org/10.1002/j.1537-2197.1987.tb08741.x.

B. Houchmandzadeh. Giant fluctuations in logistic growth of two species competing for limited resources. *Physical Review E*, 98(4), Oct. 2018. doi: 10.1103/physreve.98.042118. URL https://doi.org/10.1103/physreve.98.042118.

L. Marrec, C. Bank, and T. Bertrand. Solving the stochastic dynamics of population growth. *Ecology and Evolution*, 13(8), July 2023. ISSN 2045-7758. doi: 10.1002/ece3.10295. URL http://dx.doi.org/10.1002/ece3.10295.

N. M. Vega and J. Gore. Stochastic assembly produces heterogeneous communities in the caenorhabditis elegans intestine. *PLOS Biology*, 15(3):e2000633, Mar. 2017. doi: 10.1371/journal.pbio.2000633. URL https://doi.org/10.1371/journal.pbio.2000633.
